## Supplementary figures and images for "MECP2pedia: A Comprehensive Transcriptome Portal for MECP2 Disease Research"

### Supplemental Figures

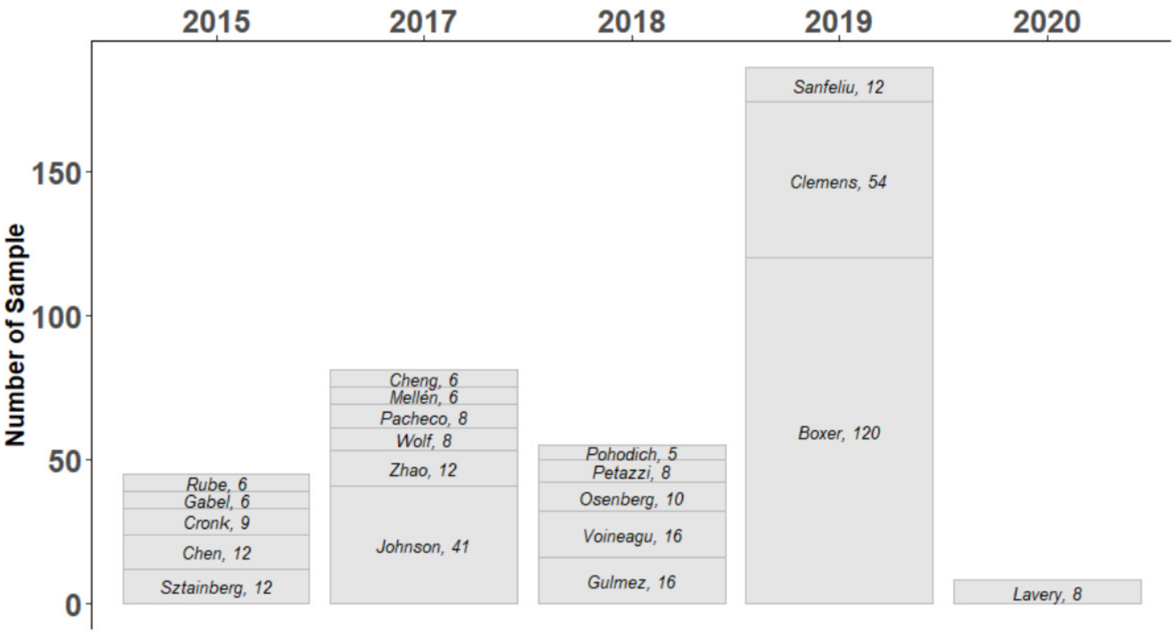

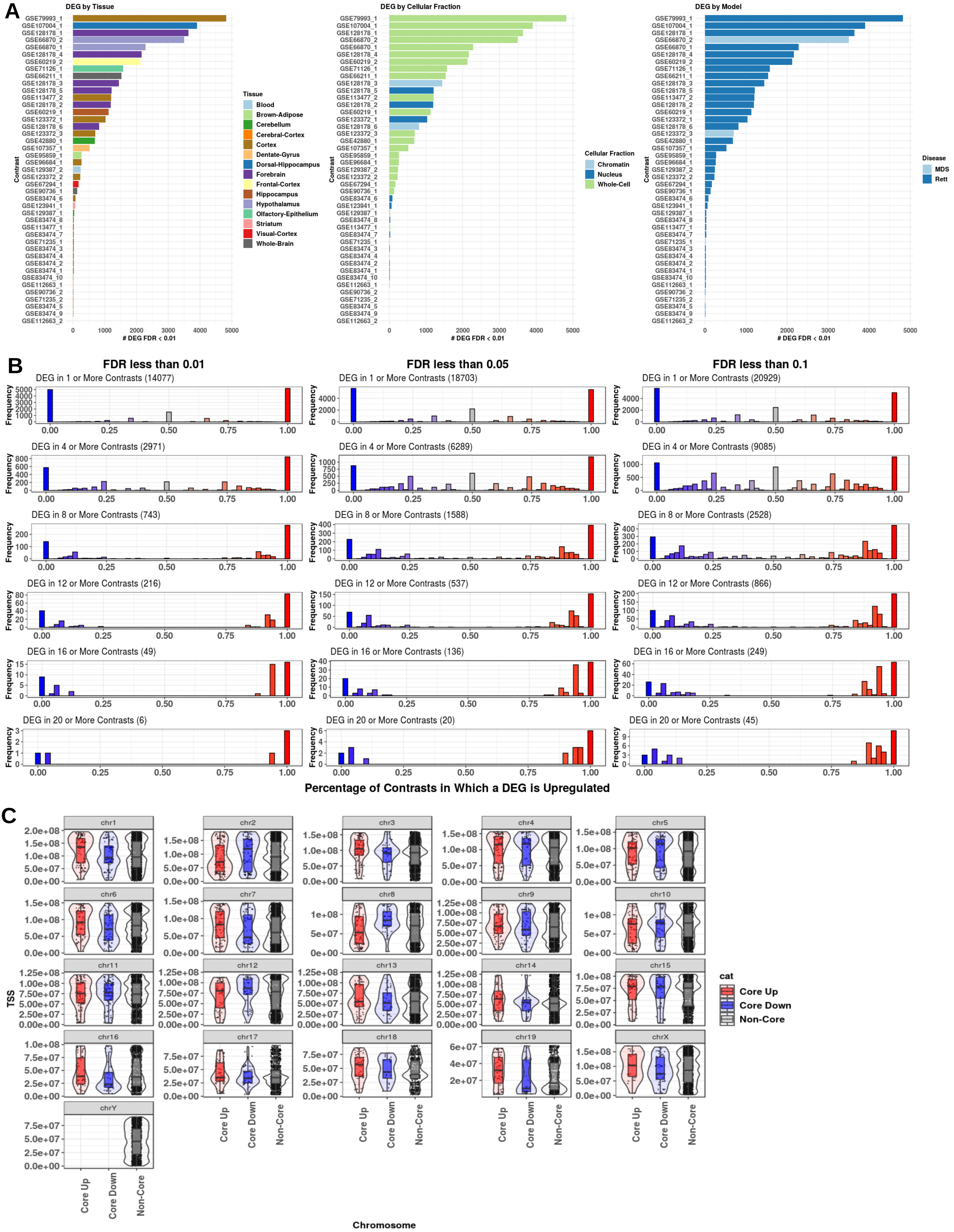

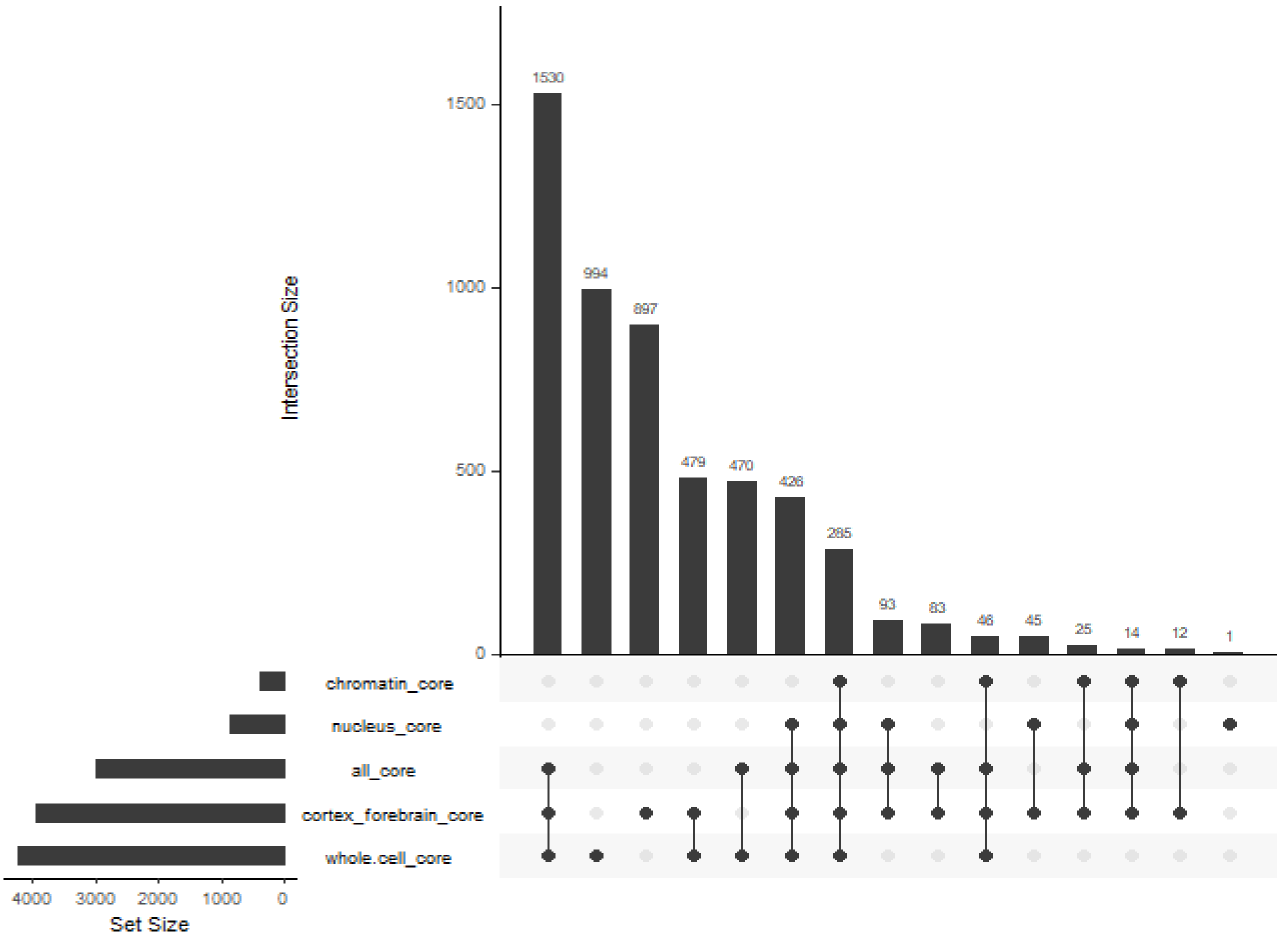

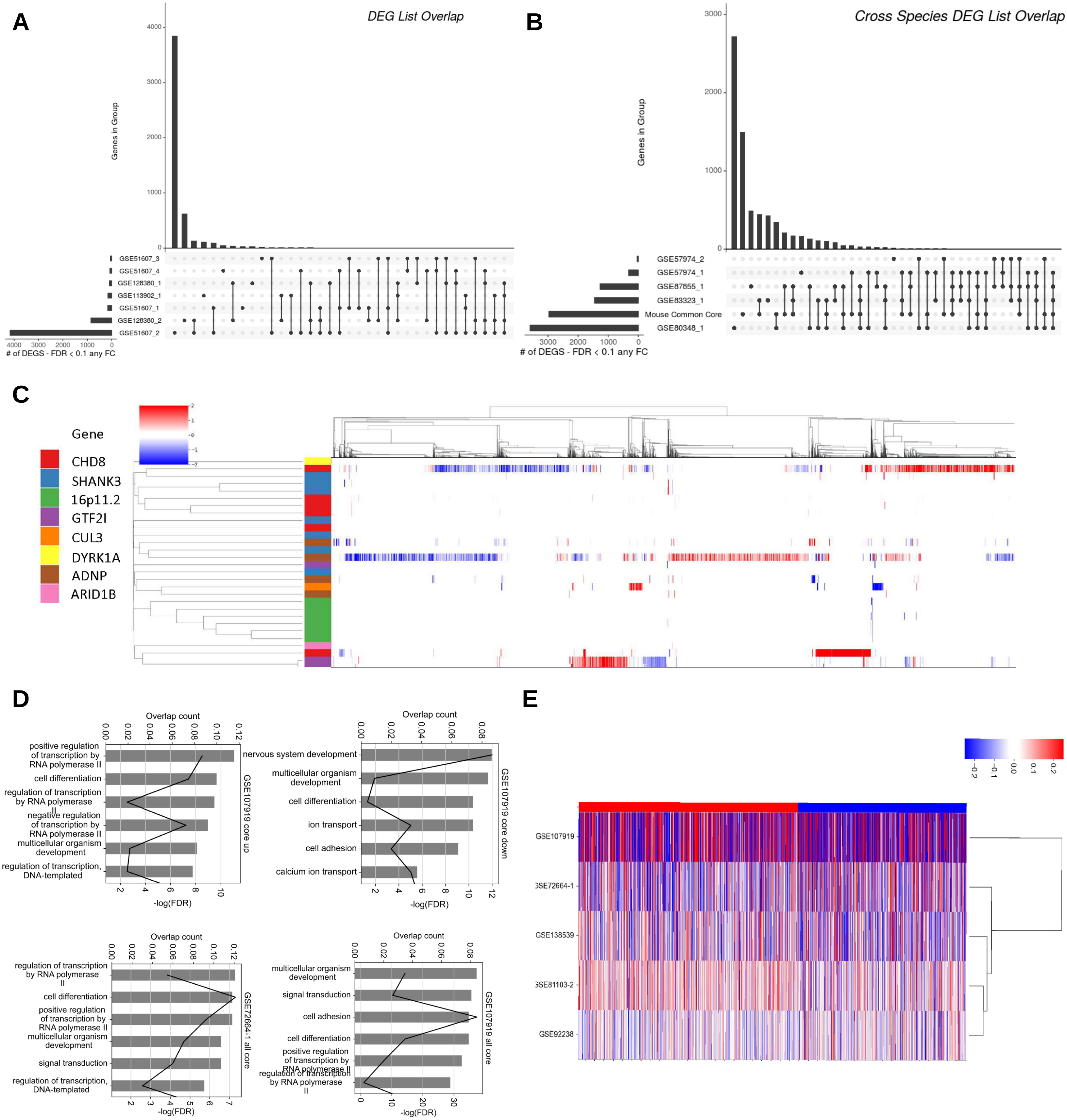

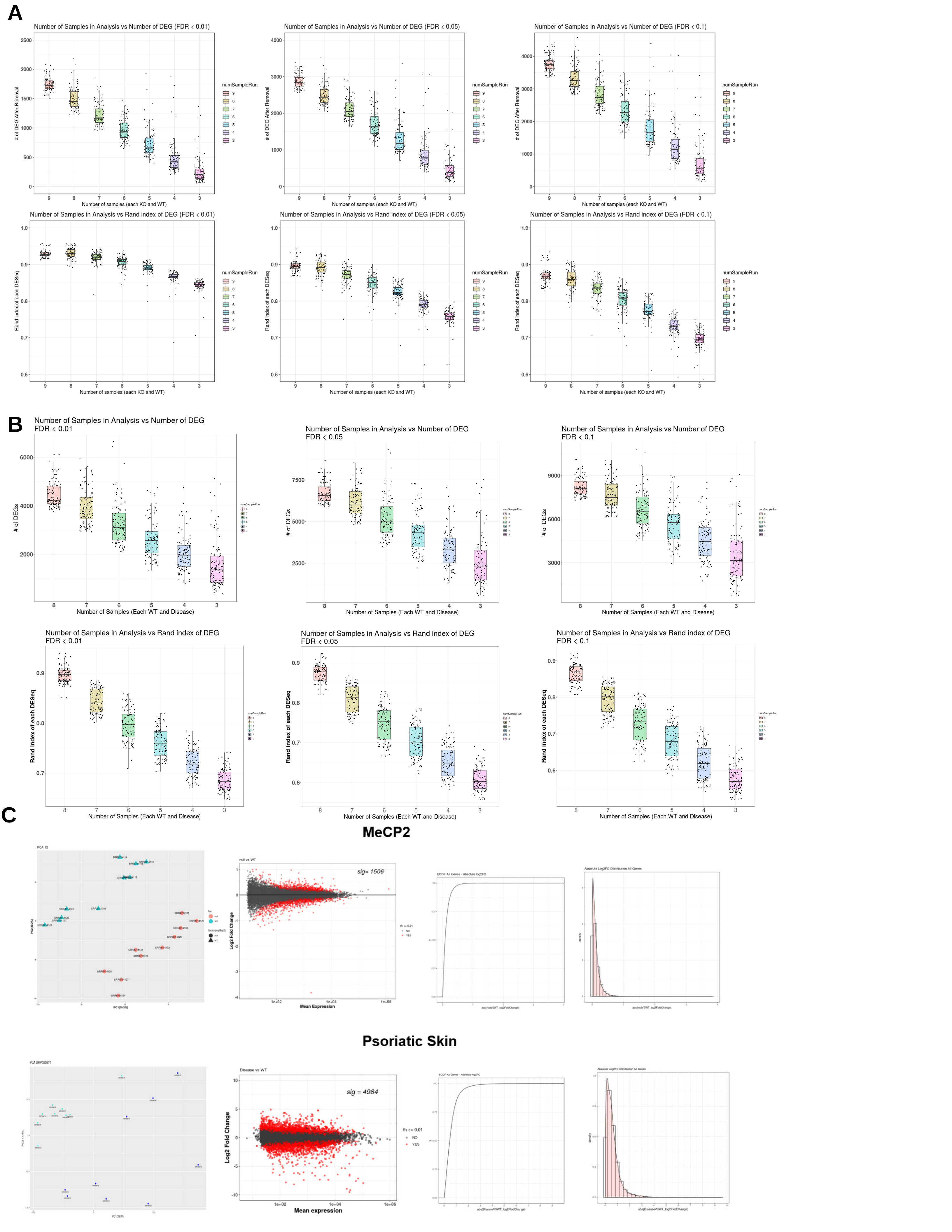

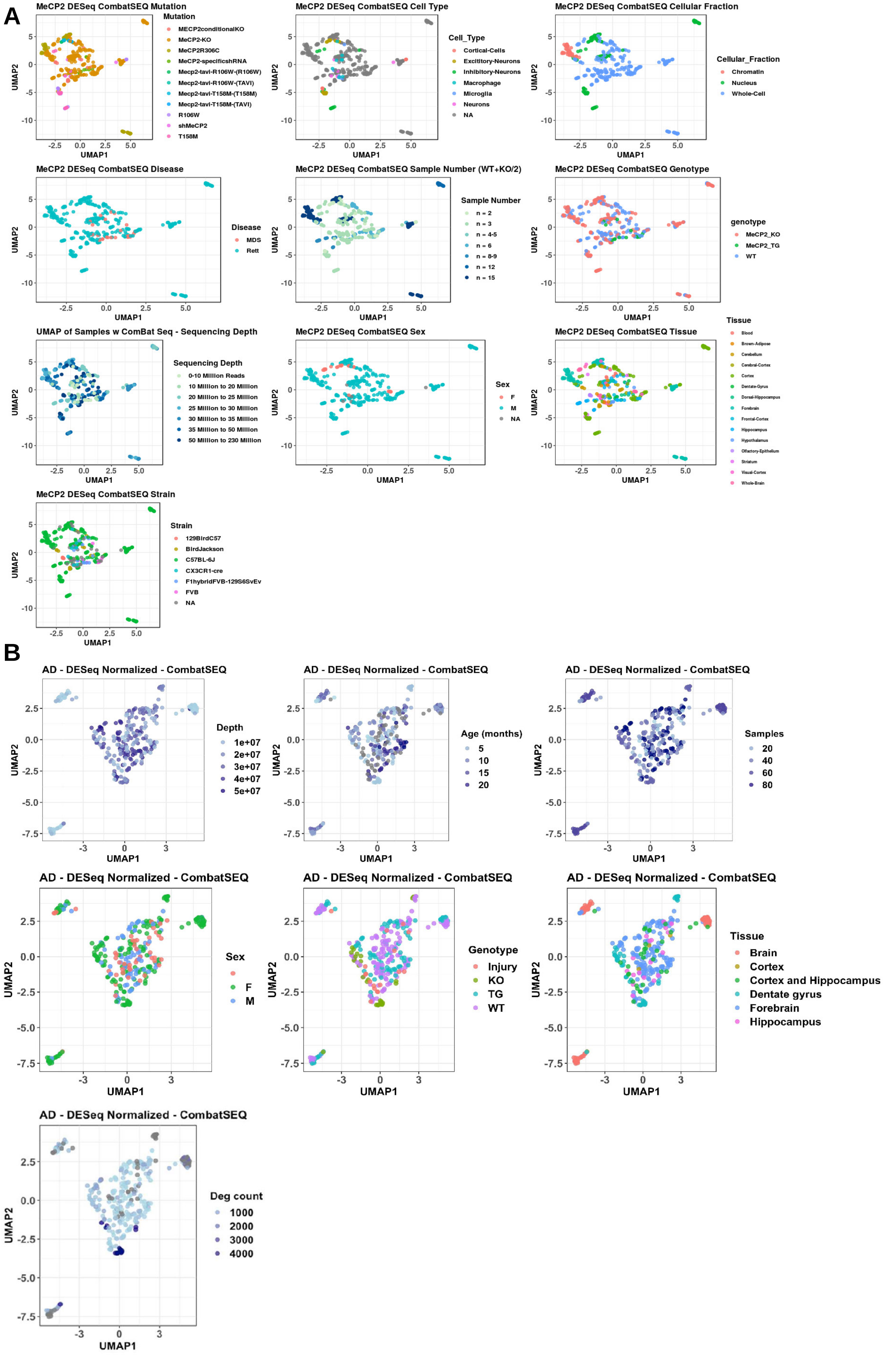
