## Supplemental Table 1 for "MECP2pedia: A Comprehensive Transcriptome Portal for MECP2 Disease Research"

| Name | GEO # | Disease | Organism | Cont.ID | Model | Sex | Age | Tissue | Strain | Cell Fraction | Mutation | Cell Type | # samples |
|---|---|---|---|---|---|---|---|---|---|---|---|---|---|
| Counts_107004_1 | GSE107004 | Rett | Mus musculus | 1 | KO | M | 36w | Dorsal-Hippocampus | C57BL-6J | Whole-Cell | MeCP2-specificshRNA | | 16 |
| Counts_107357_1 | GSE107357 | Rett | Mus musculus | 1 | KO | M | 8w | Dentate-Gyrus | FVB | Whole-Cell | MeCP2-KO | | 6 |
| Counts_112663_1 | GSE112663 | Rett | Mus musculus | 1 | KO | M | 22w | Cerebral-Cortex | BirdJackson | Whole-Cell | MECP2conditionalKO | | 8 |
| Counts_112663_2 | GSE112663 | Rett | Mus musculus | 2 | KO | M | 22w | Cerebellum | BirdJackson | Whole-Cell | MECP2conditionalKO | | 8 |
| Counts_113477_1 | GSE113477 | Rett | Mus musculus | 1 | KO | M | 7w | Hippocampus | C57BL-6J | Whole-Cell | MeCP2-KO | | 18 |
| Counts_113477_2 | GSE113477 | Rett | Mus musculus | 2 | KO | M | 7w | Cortex | C57BL-6J | Whole-Cell | MeCP2-KO | | 12 |
| Counts_123372_1 | GSE123372 | Rett | Mus musculus | 1 | KO | M | 7w | Cortex | C57BL-6J | Nucleus | MeCP2-KO | | 24 |
| Counts_123372_2 | GSE123372 | Rett | Mus musculus | 2 | KO | M | 7w | Cortex | C57BL-6J | Whole-Cell | MeCP2-KO | | 12 |
| Counts_123372_3 | GSE123372 | MDS | Mus musculus | 3 | TG | M | 7w | Cortex | FVB | Whole-Cell | MeCP2-TG | | 10 |
| Counts_123941_1 | GSE123941 | Rett | Mus musculus | 1 | KO | M | 6w | Striatum | C57BL-6J | Nucleus | MeCP2-KO | Inhibitory-Neurons | 8 |
| Counts_128178_1 | GSE128178 | Rett | Mus musculus | 1 | KO | M | 8w | Forebrain | C57BL-6J | Whole-Cell | MeCP2-KO | | 30 |
| Counts_128178_2 | GSE128178 | Rett | Mus musculus | 2 | KO | M | 8w | Forebrain | C57BL-6J | Nucleus | MeCP2-KO | | 30 |
| Counts_128178_3 | GSE128178 | Rett | Mus musculus | 3 | KO | M | 8w | Forebrain | C57BL-6J | Chromatin | MeCP2-KO | | 30 |
| Counts_128178_4 | GSE128178 | Rett | Mus musculus | 4 | KO | M | 8w | Forebrain | C57BL-6J | Whole-Cell | MeCP2R306C | | 30 |
| Counts_128178_5 | GSE128178 | Rett | Mus musculus | 5 | KO | M | 8w | Forebrain | C57BL-6J | Nucleus | MeCP2R306C | | 30 |
| Counts_128178_6 | GSE128178 | Rett | Mus musculus | 6 | KO | M | 8w | Forebrain | C57BL-6J | Chromatin | MeCP2R306C | | 30 |
| Counts_129387_1 | GSE129387 | Rett | Mus musculus | 1 | KO | M | 7w | Cerebellum | C57BL-6J | Whole-Cell | MeCP2-KO | | 6 |
| Counts_129387_2 | GSE129387 | Rett | Mus musculus | 2 | KO | M | 7w | Blood | C57BL-6J | Whole-Cell | MeCP2-KO | | 6 |
| Counts_42880_1 | GSE42880 | Rett | Mus musculus | 1 | KO | M | 7w | Cerebellum | C57BL-6J | Whole-Cell | MeCP2-KO | | 6 |
| Counts_60219_1 | GSE60219 | Rett | Mus musculus | 1 | KO | M | | Hippocampus | | Whole-Cell | MeCP2-KO | | 4 |
| Counts_60219_2 | GSE60219 | Rett | Mus musculus | 2 | KO | M | | Frontal-Cortex | | Whole-Cell | MeCP2-KO | | 4 |
| Counts_66211_1 | GSE66211 | Rett | Mus musculus | 1 | KO | M | 10w | Whole-Brain | | Whole-Cell | MeCP2-KO | Microglia | 9 |
| Counts_66870_1 | GSE66870 | Rett | Mus musculus | 1 | KO | M | 7w | Hypothalamus | | Whole-Cell | MeCP2-KO | | 6 |
| Counts_66870_2 | GSE66870 | MDS | Mus musculus | 2 | TG | M | 7w | Hypothalamus | | Whole-Cell | MeCP2-TG | | 6 |
| Counts_67294_1 | GSE67294 | Rett | Mus musculus | 1 | KO | M | 9w | Visual-Cortex | | Whole-Cell | MeCP2-KO | | 6 |
| Counts_71126_1 | GSE71126 | Rett | Mus musculus | 1 | KO | M | 8w | Olfactory-Epithelium | 129BirdC57 | Whole-Cell | MeCP2-KO | | 6 |
| Counts_71235_1 | GSE71235 | MDS | Mus musculus | 1 | TG | M | 4w | Hippocampus | F1hybridFVB-129S6SvEv | Whole-Cell | MeCP2-TG | | 6 |
| Counts_71235_2 | GSE71235 | MDS | Mus musculus | 2 | TG | M | 8w | Hippocampus | F1hybridFVB-129S6SvEv | Whole-Cell | MeCP2-TG | | 6 |
| Counts_79993_1 | GSE79993 | Rett | Mus musculus | 1 | KO | | E14.5 | Cortex | | Whole-Cell | shMeCP2 | Neurons | 6 |
| Counts_83474_10 | GSE83474 | Rett | Mus musculus | 10 | KO | F | 18w | Cortex | C57BL-6J | Nucleus | Mecp2-tavi-T158M-(T158M) | Excititory-Neurons | 4 |
| Counts_83474_1 | GSE83474 | Rett | Mus musculus | 1 | KO | M | 6w | Cortex | C57BL-6J | Nucleus | R106W | Excititory-Neurons | 8 |
| Counts_83474_2 | GSE83474 | Rett | Mus musculus | 2 | KO | M | 6w | Cortex | C57BL-6J | Nucleus | T158M | Inhibitory-Neurons | 8 |
| Counts_83474_3 | GSE83474 | Rett | Mus musculus | 3 | KO | M | 6w | Cortex | C57BL-6J | Nucleus | R106W | Cortical-Cells | 4 |
| Counts_83474_4 | GSE83474 | Rett | Mus musculus | 4 | KO | M | 6w | Cortex | C57BL-6J | Whole-Cell | R106W | Cortical-Cells | 4 |
| Counts_83474_5 | GSE83474 | Rett | Mus musculus | 5 | KO | M | 6w | Cortex | C57BL-6J | Nucleus | T158M | Excititory-Neurons | 8 |
| Counts_83474_6 | GSE83474 | Rett | Mus musculus | 6 | KO | M | 6w | Cortex | C57BL-6J | Nucleus | R106W | Inhibitory-Neurons | 8 |
| Counts_83474_7 | GSE83474 | Rett | Mus musculus | 7 | KO | F | 18w | Cortex | C57BL-6J | Nucleus | Mecp2-tavi-R106W-(TAVI) | Excititory-Neurons | 4 |
| Counts_83474_8 | GSE83474 | Rett | Mus musculus | 8 | KO | F | 18w | Cortex | C57BL-6J | Nucleus | Mecp2-tavi-R106W-(R106W) | Excititory-Neurons | 4 |
| Counts_83474_9 | GSE83474 | Rett | Mus musculus | 9 | KO | F | 18w | Cortex | C57BL-6J | Nucleus | Mecp2-tavi-T158M-(TAVI) | Excititory-Neurons | 4 |
| Counts_90736_1 | GSE90736 | Rett | Mus musculus | 1 | KO | F | 5w | Whole-Brain | C57BL-6J | Whole-Cell | MeCP2-KO | Microglia | 6 |
| Counts_90736_2 | GSE90736 | Rett | Mus musculus | 2 | KO | F | 24w | Whole-Brain | C57BL-6J | Whole-Cell | MeCP2-KO | Microglia | 6 |
| Counts_95859_1 | GSE95859 | Rett | Mus musculus | 1 | KO | M | | Brown-Adipose | CX3CR1-cre | Whole-Cell | MeCP2-KO | Macrophage | 8 |
| Counts_96684_1 | GSE96684 | Rett | Mus musculus | 1 | KO | M | 9w | Cortex | C57BL-6J | Whole-Cell | MeCP2-KO | | 8 |
