## Supplemental Table 3 for "MECP2pedia: A Comprehensive Transcriptome Portal for MECP2 Disease Research"

| Name | GEO # | Tissue | 1st author | Gene | Genotype | Strain | Sex | Age (Weeks) | # samples | FDR<0.01 |
|---|---|---|---|---|---|---|---|---|---|---|
| GSE107919 | GSE107919 | | Richard Lu | CHD8 | KO | mixed | | | 4 | 7859 |
| GSE109328-1 | GSE109328 | prefrontal cortex | Zhen Yan | SHANK3 | saline control | | | 5 | 6 | 33 |
| GSE109328-2 | GSE109328 | prefrontal cortex | Zhen Yan | SHANK3 | romidepsin treated | | | 5 | 5 | 21 |
| GSE113368 | GSE113368 | prefrontal cortex | Kihoon Han | SHANK3 | mutant | C57BL/6J | | 10 | 6 | 1 |
| GSE113754 | GSE113754 | prefrontal cortex | Lucia Peixoto | SHANK3 | mutant | | M | 8 | 20 | 16 |
| GSE124946-1 | GSE124946 | striatum | Kihoon Han | SHANK3 | KO | C57BL/6J | | 10 | 6 | 4 |
| GSE124946-2 | GSE124946 | striatum | Kihoon Han | SHANK3 | KO | C57BL/6J | | 10 | 6 | 1 |
| GSE126559-1 | GSE126559 | cerebellum | Michael Talkowski | 16P11.2 | KO | C57BL/6N:129Sv | M | | 8 | 20 |
| GSE126559-2 | GSE126559 | cerebellum | Michael Talkowski | 16P11.2 | KO | C57BL/6N:129Sv | F | | 8 | 24 |
| GSE126559-3 | GSE126559 | cortex | Michael Talkowski | 16P11.2 | KO | C57BL/6N:129Sv | M | | 8 | 23 |
| GSE126559-4 | GSE126559 | cortex | Michael Talkowski | 16P11.2 | KO | C57BL/6N:129Sv | F | | 8 | 26 |
| GSE126559-5 | GSE126559 | striatum | Michael Talkowski | 16P11.2 | KO | C57BL/6N:129Sv | M | | 8 | 24 |
| GSE126559-6 | GSE126559 | striatum | Michael Talkowski | 16P11.2 | KO | C57BL/6N:129Sv | F | | 8 | 37 |
| GSE128838Xx-xXGSE128841 | GSE128838Xx-xXGSE128841 | cortex | Boaz Barak | GTF2I | heterozygote mutation | C57BL/6J | | | 6 | 15 |
| GSE133160 | GSE133160 | | Viraj Sanghvi | CUL3 | mutant | | | | 5 | 588 |
| GSE138539 | GSE138539 | striatum | Evan Elliott | SHANK3 | KO | C57BL/6J | | 8 | 12 | 154 |
| GSE67052 | GSE67052 | | John Crispino | DYRK1A | cKO | C57BL/6J | | 4 | 10 | 6 |
| GSE72442 | GSE72442 | cortex | Li-Huei Tsai | CHD8 | KO | | | embryonic | 10 | 1520 |
| GSE72664-1 | GSE72664 | hippocampus | Metsada Pasmanik-Ch | ADNP | heterozygote mutation | mixed C57BL and 129/SvJ background | F | 4 | 5 | 1277 |
| GSE72664-2 | GSE72664 | hippocampus | Metsada Pasmanik-Ch | ADNP | heterozygote mutation | mixed C57BL and 129/SvJ background | M | 4 | 7 | 309 |
| GSE72664-3 | GSE72664 | hippocampus | Metsada Pasmanik-Ch | ADNP | heterozygote mutation | mixed C57BL and 129/SvJ background | F | 20 | 6 | 9262 |
| GSE72664-4 | GSE72664 | hippocampus | Metsada Pasmanik-Ch | ADNP | heterozygote mutation | mixed C57BL and 129/SvJ background | M | 20 | 5 | 39 |
| GSE80803Xx-xXGSE87370 | GSE80803Xx-xXGSE87370 | whole brain | Eunjoon Kim | CHD8 | heterozygote mutation | C57BL/6J | | 0 | 12 | 1 |
| GSE81082 | GSE81082 | lip | Susan Corley | GTF2I | KO | C57BL/6J | | | 6 | 2281 |
| GSE81103-1 | GSE81103 | cortex | Albert Basson | CHD8 | heterozygote mutation | C57BL/6J | | embryonic | 6 | 12 |
| GSE81103-2 | GSE81103 | cortex | Albert Basson | CHD8 | heterozygote mutation | C57BL/6J | | 1 | 6 | 649 |
| GSE87370Xx-xXGSE103377 | GSE87370Xx-xXGSE103377 | hippocampus | Eunjoon Kim | CHD8 | heterozygote mutation | C57BL/6J | | 4 | 12 | 45 |
| GSE92238 | GSE92238 | hippocampus | Hao Zhu | ARID1B | heterozygote mutation | C57/B6 | M | 11 | 8 | 60 |
| GSE97471 | GSE97471 | lip | Susan Corley | GTF2I | Gtf2ird1tm1Hrd | | | | 6 | 2319 |
