## Supplemental Table 5 for "MECP2pedia: A Comprehensive Transcriptome Portal for MECP2 Disease Research"

| First Author | Names | GEO # | Disease | Organism | Cont.ID | Sex | Age | Tissue | Cell_line | Strain | cellular_fraction | Mutation | Cell_type |
|---|---|---|---|---|---|---|---|---|---|---|---|---|---|
| Renthal | Counts_113902_1_FDR | GSE113902 | Rett | Homo sapiens | 1 | F | NA | Occipital-Cortex | NA | NA | Nucleus | Mosaic | NA |
| Aldinger | Counts_128380_1_FDR | GSE128380 | Rett | Homo sapiens | 1 | F | NA | Cingulate-Cortex | NA | NA | Whole-Cell | NA | NA |
| Aldinger | Counts_128380_2_FDR | GSE128380 | Rett | Homo sapiens | 2 | F | NA | Temporal-Cortex | NA | NA | Whole-Cell | NA | NA |
| Tanaka | Counts_51607_1_FDR | GSE51607 | Rett | Homo sapiens | 1 | NA | NA | NA | iPSC | NA | Whole-Cell | T158M | iPSC |
| Tanaka | Counts_51607_2_FDR | GSE51607 | Rett | Homo sapiens | 2 | NA | NA | NA | iPSC | NA | Whole-Cell | X487W | iPSC |
| Tanaka | Counts_51607_3_FDR | GSE51607 | Rett | Homo sapiens | 3 | NA | NA | NA | iPSC | NA | Whole-Cell | R306C | iPSC |
| Tanaka | Counts_51607_4_FDR | GSE51607 | Rett | Homo sapiens | 4 | NA | NA | NA | iPSC | NA | Whole-Cell | E235fs | iPSC |
| Liu | Counts_57974_1_FDR | GSE57974 | Rett | Macaca fascicularis | 1 | NA | P0 | Brain | NA | NA | Whole-Cell | NA | NA |
| Liu | Counts_57974_2_FDR | GSE57974 | MDS | Macaca fascicularis | 2 | NA | P0 | Brain | NA | NA | Whole-Cell | NA | NA |
| Veeraragavan | Counts_83323_1_FDR | GSE83323 | Rett | Rattus norvegicus | 1 | M | 6w | Hypothalamus | NA | NA | Whole-Cell | NA | NA |
| Bhattacherjee | Counts_87855_1_FDR | GSE87855 | Rett | Rattus norvegicus | 1 | M | 4w | Dorsal Root Ganglia | NA | NA | Whole-Cell | NA | Dorsal Root Ganglia |
| Van der Vaart | Counts_80348_1_FDR | GSE80348 | Rett | Danio rerio | 1 | M | 6h after fertilization | Whole Embryo | NA | NA | Whole-Cell | NA | NA |
