## Supplemental Table 6 for "MECP2pedia: A Comprehensive Transcriptome Portal for MECP2 Disease Research"

| Name | Enrichment to Core-Down Genes | | | | | | | | Enrichment to Core-Up Genes | | | | | | | |
|---|---|---|---|---|---|---|---|---|---|---|---|---|---|---|---|---|
| | SIZE | ES | NES | NOM.p.val | FDR.q.val | FWER.p.val | RANK.AT.MAX | LEADING.EDGE | SIZE | ES | NES | NOM.p.val | FDR.q.val | FWER.p.val | RANK.AT.MAX | LEADING.EDGE |
| GSE113902_1 | 1015 | 0.921258 | 0.971319 | 0.898 | 0.935335 | 1 | 317 | tags=5%, list=2%, signal=4% | 1401 | 0.958753 | 1.009798 | 0.346 | 1 | 0.706 | 373 | tags=6%, list=3%, signal=6% |
| GSE128380_1 | 1055 | -0.59527 | -1.59824 | 0.003976143 | 0.003452 | 0.002 | 1473 | tags=25%, list=11%, signal=26% | 1456 | -0.36277 | -0.97948 | 0.40816328 | 0.428654 | 0.516 | 1233 | tags=14%, list=9%, signal=14% |
| GSE128380_2 | 1041 | -0.46299 | -1.47374 | 0 | 0 | 0 | 1601 | tags=24%, list=12%, signal=25% | 1442 | 0.375051 | 1.026305 | 0.38951695 | 0.387167 | 0.757 | 1413 | tags=16%, list=11%, signal=16% |
| GSE51607_1 | 1006 | 0.445294 | 0.884388 | 0.79094076 | 0.883452 | 0.513 | 558 | tags=5%, list=4%, signal=5% | 1404 | 0.477342 | 0.979128 | 0.53556484 | 0.791815 | 0.332 | 589 | tags=6%, list=4%, signal=6% |
| GSE51607_2 | 954 | -0.42768 | -1.41146 | 0 | 0 | 0 | 2866 | tags=30%, list=22%, signal=36% | 1377 | -0.25718 | -0.85792 | 0.9569138 | 0.965167 | 1 | 2124 | tags=15%, list=16%, signal=16% |
| GSE51607_3 | 1004 | -0.80176 | -0.90185 | 0.81171066 | 0.821978 | 0.973 | 104 | tags=1%, list=1%, signal=1% | 1401 | -0.9586 | -1.08574 | 0.2862109 | 0.915832 | 0.593 | 105 | tags=2%, list=1%, signal=2% |
| GSE51607_4 | 957 | 0.586899 | 0.948996 | 0.47308782 | 0.494784 | 0.423 | 338 | tags=5%, list=3%, signal=4% | 1346 | 0.592684 | 0.967576 | 0.4770115 | 0.707326 | 0.407 | 295 | tags=5%, list=2%, signal=5% |
| GSE57974_1 | 1079 | -0.5302 | -1.39173 | 0.004110997 | 0.0015 | 0.004 | 2168 | tags=22%, list=15%, signal=24% | 1481 | -0.57074 | -1.51582 | 0 | 0 | 0 | 2240 | tags=24%, list=16%, signal=26% |
| GSE57974_2 | 1079 | 0.926797 | 2.04814 | 0.15766738 | 0.105777 | 0.073 | 24 | tags=0%, list=0%, signal=0% | 1481 | -0.49674 | -1.3061 | 0.15900382 | 0.119048 | 0.175 | 2266 | tags=25%, list=16%, signal=27% |
| GSE80348_1 | 575 | -0.54906 | -1.0792 | 0.29297298 | 0.803303 | 0.574 | 1026 | tags=14%, list=11%, signal=15% | 855 | -0.37707 | -0.74537 | 0.9829787 | 0.982316 | 0.999 | 952 | tags=10%, list=10%, signal=10% |
| GSE83323_1 | 1076 | -0.75239 | -1.75399 | 0 | 0 | 0 | 983 | tags=27%, list=7%, signal=27% | 1437 | 0.79153 | 1.9779 | 0 | 0 | 0 | 870 | tags=29%, list=6%, signal=28% |
| GSE87855_1 | 1002 | -0.65003 | -1.30348 | 0.03002611 | 0.033904 | 0.043 | 663 | tags=16%, list=5%, signal=15% | 1376 | 0.534918 | 1.236552 | 0.023391813 | 0.032321 | 0.016 | 992 | tags=22%, list=8%, signal=21% |
